# The molecular chaperone TRAP1 promotes translation of *Luc7I3* mRNA to enhance ovarian cancer cell proliferation

**DOI:** 10.1101/2025.04.01.646636

**Authors:** Sabrina De Lella, Lorenza Pedalino, Mehad Almagboul Abdalla Abaker, Chiara Mignogna, Raffaele Cautiero, Franca Esposito, Danilo Swann Matassa, Rosario Avolio

**Author notes:** Correspondence may also be addressed to: Danilo Swann Matassa.

## Abstract

Heat shock proteins have been increasingly identified in RNA-interactomes, suggesting potential roles beyond their canonical functions. Among those, the cancer-linked chaperone TRAP1 has been mainly characterized for its regulatory role on respiratory complex activity and protein synthesis, while its specific function as an RNA-binding protein (RBP) remains unclear. In this study, we confirmed the RNA-binding activity of TRAP1 in living cells using both protein- and RNA-centric approaches and demonstrated that multiple TRAP1 regions cooperate in such binding. Enhanced cross-linking and immunoprecipitation (eCLIP) in high-grade serous ovarian cancer cells revealed that TRAP1 primarily binds cytosolic protein-coding genes, with the majority coding for splicing-related factors. Notably, among TRAP1 most significantly bound transcripts, we identified the splicing factor LUC7L3, a U1 snRNP component involved in cell proliferation. We confirmed TRAP1 binding to *Luc7l3* transcript by RIP-qPCR and showed that TRAP1 promotes *Luc7l3* mRNA translation. Furthermore, we demonstrated that TRAP1 enhances ovarian cancer cell proliferation through LUC7L3 translational regulation. In summary, our findings provide the first comprehensive characterization of TRAP1 as an RBP and identify a critical target for ovarian cancer cell proliferation, offering new insights into its multifaceted roles in tumor biology.

## INTRODUCTION

RNA-binding proteins (RBPs) are critical regulators of gene expression (Gebauer et al., 2021). Together with messenger RNA (mRNA) and non-coding RNA they take part in the assembly of ribonucleoprotein complexes soon after transcription and are involved in all steps of RNA metabolism (Glisovic et al., 2008). The family of RBPs comprises a vast group of diverse members, whose number has dramatically expanded within the last two decades with the advent of transcriptome-wide RNA-interactome approaches aiming at isolating the RBP-repertoire (named RBPome) of cells and tissues (Perez-Perri et al., 2023; Steinmetz et al., 2023). The identification of hundreds of novel RNA-binders lacking known RNA-binding domains (RBDs) led to the classification of these proteins as “non-canonical” RBPs (Hentze et al., 2018).

Non-canonical RBPs include proteins with previously established functions not related with RNA. Metabolic enzymes are one such example (Castello et al., 2015) where the RNA targets and possible roles in post-transcriptional gene regulation of several members of this family have been addressed (Chu et al., 1991; Hentze & Argos, 1991; Kejiou et al., 2023; Noble et al., 2024; Zhou et al., 2008). At the same time, the metabolic activity of enzymes is regulated by RNA, introducing the concept of riboregulation (Guiducci et al., 2019; Huppertz et al., 2022). Together with metabolic enzymes, heat shock proteins (HSPs) have been consistently detected as RBPs by RNA-interactome approaches (Moore et al., 2018). These molecular chaperones exert cytoprotective functions under stress conditions and play essential roles in protein synthesis and degradation (Hu et al., 2022). Yet, their function as RBPs has been poorly explored.

The cancer-linked chaperone TRAP1 (TNF-receptor associated protein 1 or HSP75) is one of the most frequently identified members of the HSP family in RNA-interactomes, either based on poly(A) or total RNA selection (Castello et al., 2012; Mullari et al., 2017a; Perez-Perri et al., 2018; Queiroz et al., 2019a; Trendel et al., 2019; Urdaneta et al., 2019). TRAP1 was initially identified as a TNF-receptor-associated protein (Chen et al., 1996) and as a chaperone for retinoblastoma during mitosis and after heat shock (Chen et al., 1996). TRAP1 was described as a chaperone able to protect cells from oxidative stress (Gesualdi et al., 2007) by inhibiting the opening of the mitochondrial permeability transition pore (Kang et al., 2007) and has been long considered the only mitochondrial HSP member. Interest towards TRAP1 has recently increased due to its contextual effects on cancer onset and progression, favoring the oncogenic phenotype in tumors relying on glycolysis while being negatively selected in tumors with predominant oxidative metabolism (Matassa et al., 2018). Although TRAP1 influence on cell metabolism still represents a puzzling scenario, evidence showed that this protein allows cancer cells to rapidly adapt to microenvironmental cues and energy demand (Wengert et al., 2022). Indeed, TRAP1 has emerged as a modulator of mitochondrial respiration through its regulatory effects on the activity of complexes I, III, IV of the electron transport chain (ETC) and F-ATP synthase (Cannino et al., 2022; Masgras et al., 2021; Matassa et al., 2022). Interestingly, TRAP1 effect on ETC complexes is not limited to regulating their activity. Further interest in TRAP1 arose from its subcellular compartmentalization, which allowed us to demonstrate that it behaves as a translation regulator both in the cytosol and mitochondria (Avolio et al., 2023). Through its binding to translation elongation factor eEF1A1 and ribosomes within the cytosolic translation apparatus, TRAP1 reduces the translation of nuclear-encoded ETC components, thus favoring their co-translational import to mitochondria (Avolio et al., 2023). Accordingly, TRAP1 binds the mitochondrial import channel TOMM40 and stimulates the coupled synthesis of mtDNA-encoded components of the respiratory complexes, potentially assisting their co-translational assembly (Avolio et al., 2023). Whether TRAP1 regulation of ETC component translation requires engagement of their transcripts, and if such role can be extended to other selective mRNA substrates, needs further clarification.

In this work, we provide the first demonstration of TRAP1 RNA-binding activity *in vivo* both in the cytosol and the mitochondria. We identify the ribosomal S5-D2 like fold domain as the main, but not solely, responsible for TRAP1 RNA-binding capacity, supporting its role as non-canonical RBP. Notably, using enhanced crosslinking and immunoprecipitation (eCLIP) we identify hundreds of cytosolic transcripts bound by TRAP1, with the vast majority coding for splicing-related factors. Among other transcripts, we show that TRAP1 binds the splicing factor LUC7L3 and demonstrate that TRAP1-mediated translation regulation of this factor leads to increased cell proliferation in high-grade serous ovarian cancer (HGSOC) cells.

## RESULTS

### The molecular chaperone TRAP1 binds RNA in a regulated fashion in human cells

TRAP1 has been detected in the RBPomes of several cell lines, but its role as RBP has never been validated. We confirmed TRAP1 RNA-binding using two orthogonal approaches in HeLa cells. First, we performed CLIP using an infrared dye to label TRAP1-RNA complexes. Briefly, HeLa TRAP1-Flag-HA cells were exposed to ultraviolet (UV) light (254 nm) to crosslink TRAP1 to target RNAs, and TRAP1-Flag-HA was immunoprecipitated from cell extracts treated with either low or high RNAse I concentrations. Following IP, the RNA was dephosphorylated, and a DNA oligonucleotide conjugated with an infrared dye was ligated at the 3’-end, allowing the detection of TRAP1-bound RNAs upon denaturing gel electrophoresis. As expected, high RNase I concentrations allowed visualization of TRAP1-RNA complexes just above the expected molecular mass of TRAP1 (Figure 1A). Specificity of the signal was assessed by carrying out the same experiment on a non-crosslinked sample. As a further orthogonal approach, we used RNA-interactome capture (RNA-IC) (Castello et al., 2013). In brief, following UV crosslinking of HeLa cells expressing TRAP1-eGFP, polyadenylated RNAs were isolated using oligo(dT) magnetic beads, and the fluorescence of both the eluates and inputs was measured to assess the enrichment of the eGFP-fusion protein. As expected, we detected significant green fluorescence in the eluates from cells expressing TRAP1-eGFP, while no signal was observed in the negative control (eGFP alone) (Figure 1B). We further confirmed RNA-binding of endogenous TRAP1 in the HGSOC cell line PEA1 using a CLIP variant that utilizes a biotin-conjugated linker, allowing TRAP1-RNA complexes visualization by streptavidin-HRP chemiluminescence reaction (Figure 1C). Finally, considering the well-established roles of TRAP1 in cell metabolism or mRNA translation (Avolio et al., 2023; Matassa et al., 2016), we evaluated TRAP1 binding to RNA upon stimuli modulating these processes. To this aim, we challenged HeLa TRAP1-Flag-HA cells with several treatments. We used Antimycin A to inhibit complex I activity and Emetine to inhibit mRNA translation; moreover, we cultured cells in either glucose or glutamine-deprived media. After these treatments, we assessed RNA binding by Orthogonal Organic Phase Separation (OOPS) (Queiroz et al., 2019b) followed by WB analysis. OOPS is based on organic phase separation, which retains RNA-RBP adducts at the interphase following *in vivo* UV crosslinking. Isolation of the interface and treatment with RNases leads to the release of RBPs in the organic phase upon a final round of phase separation. Our results showed that treatment with Antimycin A and glucose deprivation slightly reduce TRAP1-RNA binding capacity, while treatment with Emetine and glutamine deprivation strongly impair TRAP1 binding to RNAs (Figure 1D). The binding to RNAs of the canonical RBP DDX6, used as a positive control, was not affected, while the negative control histone H3 did not bind RNA in any condition, thus confirming the specificity of these effects. Taken together, our results provide the first demonstration that TRAP1 is a *bona fide* RNA-binding protein, and that RNA-binding is regulated during cell metabolism and translation.

**Figure 1:**
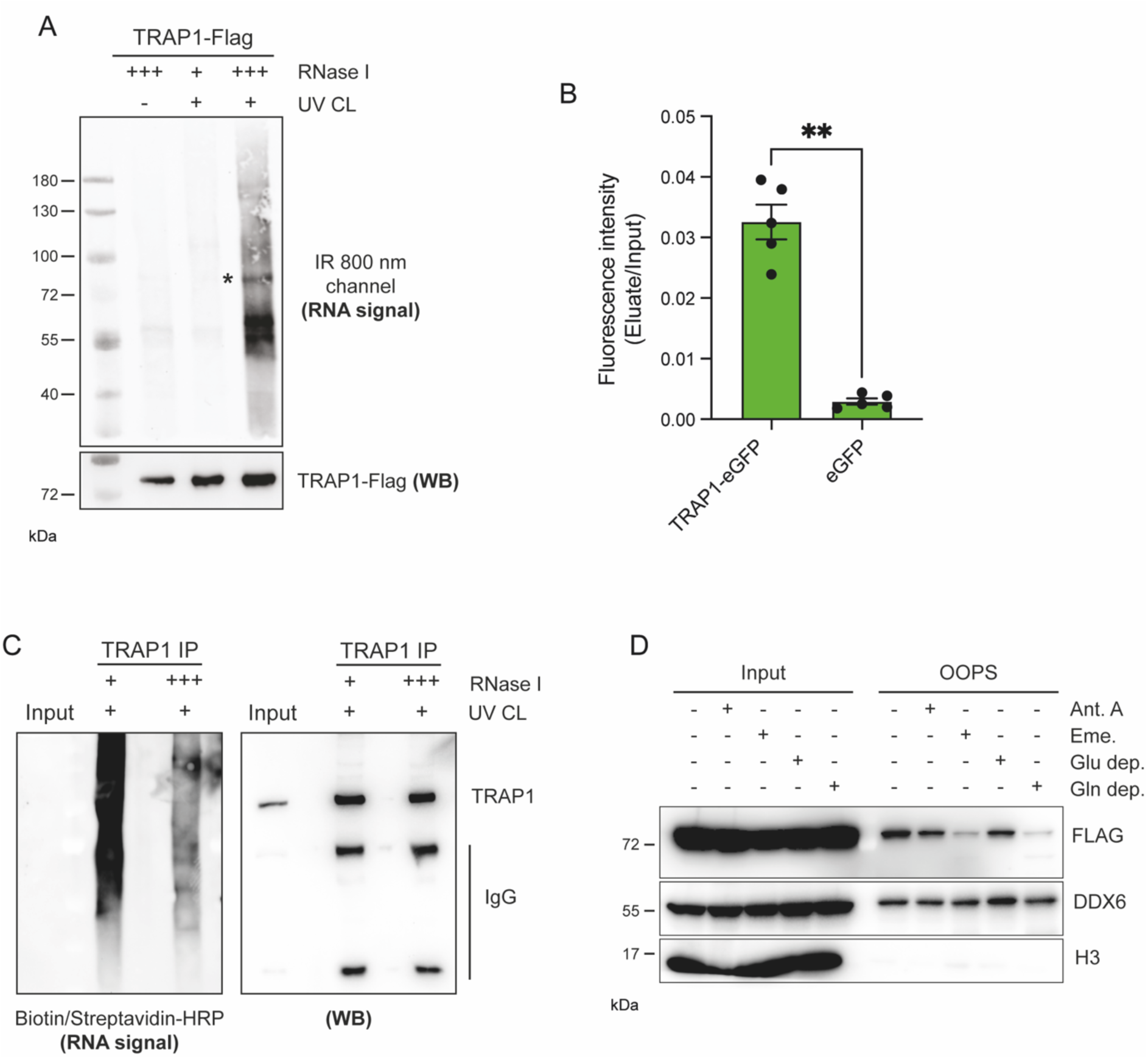
TRAP1 binds RNA in living cells. (**A**) RNA-binding activity of TRAP1-Flag-HA in HeLa FITR cells. RNA copurified with TRAP1 was dephosphorylated and ligated at the 3’-end to an infrared preA-L3-IR800-biotin DNA adaptor in order to be visualized at 800 nm (top panel). Representative western blot to confirm equal IP efficiencies (bottom panel). Asterisk indicates the expected molecular mass of TRAP1. (**B**) eGFP-based RNA-binding assay. Relative green fluorescence signal of RNA-bound fraction over input from cells expressing either TRAP1-eGFP or eGFP proteins (n = 5). Significance was assessed by unpaired Student’s t test (*p < 0.05, **p < 0.01, and ***p < 0.001). Error bars represent SEM. (**C**) RNA-binding activity of TRAP1 in PEA1 cells. Protein-RNA complexes are visualized by ligation of a biotinylated oligo to bound RNA and then detected with an HRP-linked streptavidin (left panel). Representative western blot to confirm equal IP efficiencies (right panel). (**D**) Assessment of RNA-binding by OOPS. HeLa TRAP1-Flag-HA cells were treated with either Antimycin A 1 μM (4 hrs) or Emetine 100 μg/mL (15 min) or grown in media without glucose or glutamine for 4 hrs. At the end of treatment, cells were UV crosslinked and subjected to OOPS, and both inputs and OOPS eluates were analyzed by western blot. TIAR and H3 were used as positive and negative controls respectively.

### TRAP1 binding to RNA requires multiple regions

Having established that TRAP1 binds cellular RNAs, we sought to explore which TRAP1 domains are responsible for such activity, considering the lack of canonical RBDs within its sequence. To this aim, we examined proteomic datasets that report putative RNA-binding regions of identified RBPs (Castello et al., 2016; He et al., 2016; Mullari et al., 2017b; Queiroz et al., 2019b). High-throughput studies, including RBR-ID, pCLAP, and OOPS, assigned six different RNA-binding peptides to TRAP1 in various regions (Figure 2A). Therefore, we generated six deletion mutants, each lacking one of the identified peptides, and tested their ability to bind RNA by OOPS in HeLa eGFP cells. As shown in Figure 2B, none of the tested mutants loses its ability to bind RNA, since they are all detected in the OOPS output as the positive control hnRNPC. To gain further insights into TRAP1 modality of RNA-binding, we interrogated the InterPro database to search for the main domains identified within TRAP1 structure. This analysis distinguished three main domains, excluding the N-terminal mitochondria targeting sequence (MTS) responsible for TRAP1 translocation inside mitochondria. These are: the ATPase domain, towards the N-terminal side responsible for TRAP1 chaperone activity; a Ribosomal protein S5 domain 2-like (S5-D2) domain, and a C-terminal dimerization domain, showing high homology with HSP90 (Figure 2A). We then expressed each TRAP1 domain fused to Flag-HA-tag in HeLa eGFP cells and assessed their binding to RNA using OOPS. Each fragment was expressed either inside or outside mitochondria by facultative inclusion of the TRAP1-MTS sequence. The results revealed that the S5-D2 domain retains most of the RNA-binding capacity of TRAP1 in both the cytosol and mitochondria (Figure 2C-D). Interestingly, the ATPase domain was also enriched in the OOPS output, although to a lesser extent (Figure 2C-D). These data indicate that the S5-D2 domain of TRAP1 is responsible for most RNA-binding activity, although optimal binding to RNA might involve cooperation between multiple regions.

**Figure 2:**
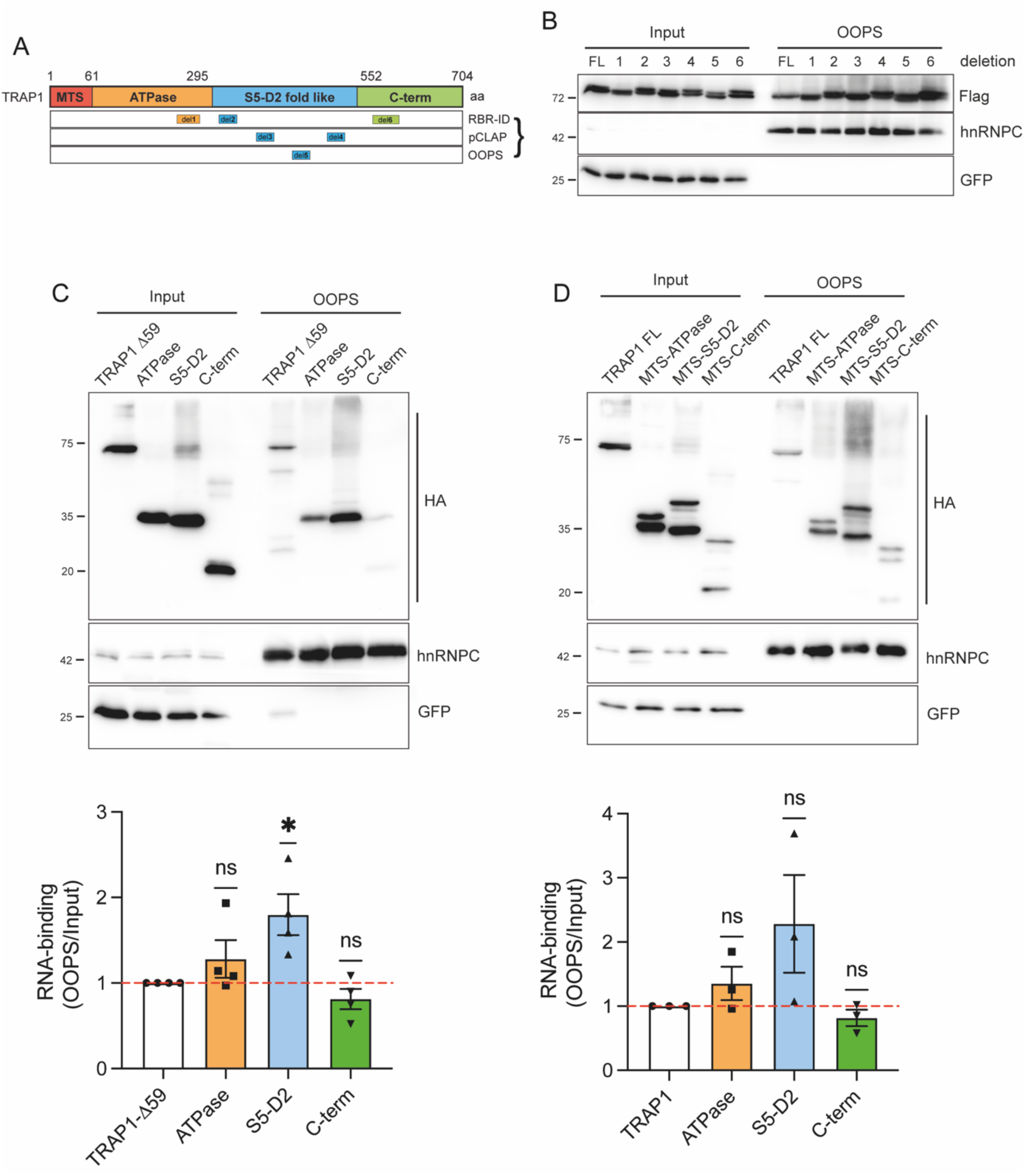
TRAP1 does not bind RNA through a well-defined domain. (**A**) Schematic representation of TRAP1 domains (ATPase, S5-D2 fold like and C-term). Peptides retrieved from RBR-ID, pCLAP and OOPS databases are indicated. MTS, mitochondria targeting sequence. (**B**) Assessment of RNA-binding of TRAP1-Flag-HA deletion mutants in HeLa FITR eGFP cells by OOPS. hnRNPC and eGFP were used as positive and negative controls respectively. (**C-D**) Assessment of RNA-binding of TRAP1 domains in HeLa FITR eGFP cells by OOPS. Left, RNA-binding analysis of domains without the MTS sequence (n = 4). Right, RNA-binding analysis of domains including the MTS sequence (n = 3). hnRNPC and eGFP were used as positive and negative controls respectively. Significance was assessed by one-sample t test. Statistical significance is represented as follows: *p < 0.05, **p < 0.01, and ***p < 0.001. Error bars represent SEM.

### TRAP1 binds to cytosolic transcripts

To identify targets binding to TRAP1, we applied eCLIP to HGSOC PEA1 cells. Our eCLIP analyses identified about five thousand TRAP1-binding regions, with 787 transcripts containing at least one significantly enriched peak (Supplementary Table 1). Interestingly, TRAP1 only binds nuclear genome-encoded mRNAs, suggesting that its functional role as RBP is achieved outside mitochondria. We found that most TRAP1 binding to its RNA substrates occurs within the CDS (Figure 3A). More precisely, a metagene analysis revealed that most TRAP1 is mainly bound near its target mRNAs’ 5’-UTR region of the CDS (Figure 3B). We also looked for enriched binding motifs among TRAP1 binding sites. We found that TRAP1 preferentially binds to purine-rich “AAG” and “GGA” stretches (Figure 3C), similarly to what has been recently reported for the homolog HSP90 (Jin et al., 2023a). Notably, we performed a pathway enrichment analysis and found that TRAP1-significantly bound RNAs are involved in protein processing at the endoplasmic reticulum (*Hsp90aa1, Hsp90ab1, Hsp90b1, Hspa5, Hspa8*) and in ribosome biogenesis (*Nop56, Tcof1, Rmb28*) (Figure 3D-E), suggesting that TRAP1 roles in these processes (Amoroso et al., 2012; Bruno et al., 2022) could at least partially involve its RNA-binding capability. Yet, to our surprise, the most enriched pathway-related term was the spliceosome (Figure 3D). Indeed, analyses of biological processes showed splicing-related terms among the most enriched ones (Supplementary Figure 1A), while analysis of cellular component showed enrichment in the nuclear lumen, nucleolus, and U2-type spliceosomal complex, among others (Supplementary Figure 1B), opening an unexplored scenario on the role of TRAP1 as RBP. Starting from these observations, we focused on the most significant TRAP1 RNA ligands. Among them, we selected LUC7L3, the human homolog of yeast U1 small nuclear RNA (snRNA)-related splicing factor Luc7p, whose functional role was recently reported (Daniels et al., 2021a). Notably, this protein was proposed to correlate with prognosis-related alternative splicing events in ovarian carcinoma (Ouyang et al., 2021). We performed RNA immunoprecipitation (RIP) followed by qPCR to validate TRAP1 binding to the *Luc7l3* transcript. Vertebrates possess two other Luc7p paralogs (LUC7L and LUC7L2), which we amplified from TRAP1 IP as controls. Our RIP-qPCR analysis confirmed a significant enrichment of *Luc7l3* mRNA in TRAP1 IP compared to the IgG control, while no enrichment was observed for either *Luc7l* or *Luc7l2*, confirming the specificity of TRAP1-*Luc7l3* binding (Figure 3F).

**Figure 3:**
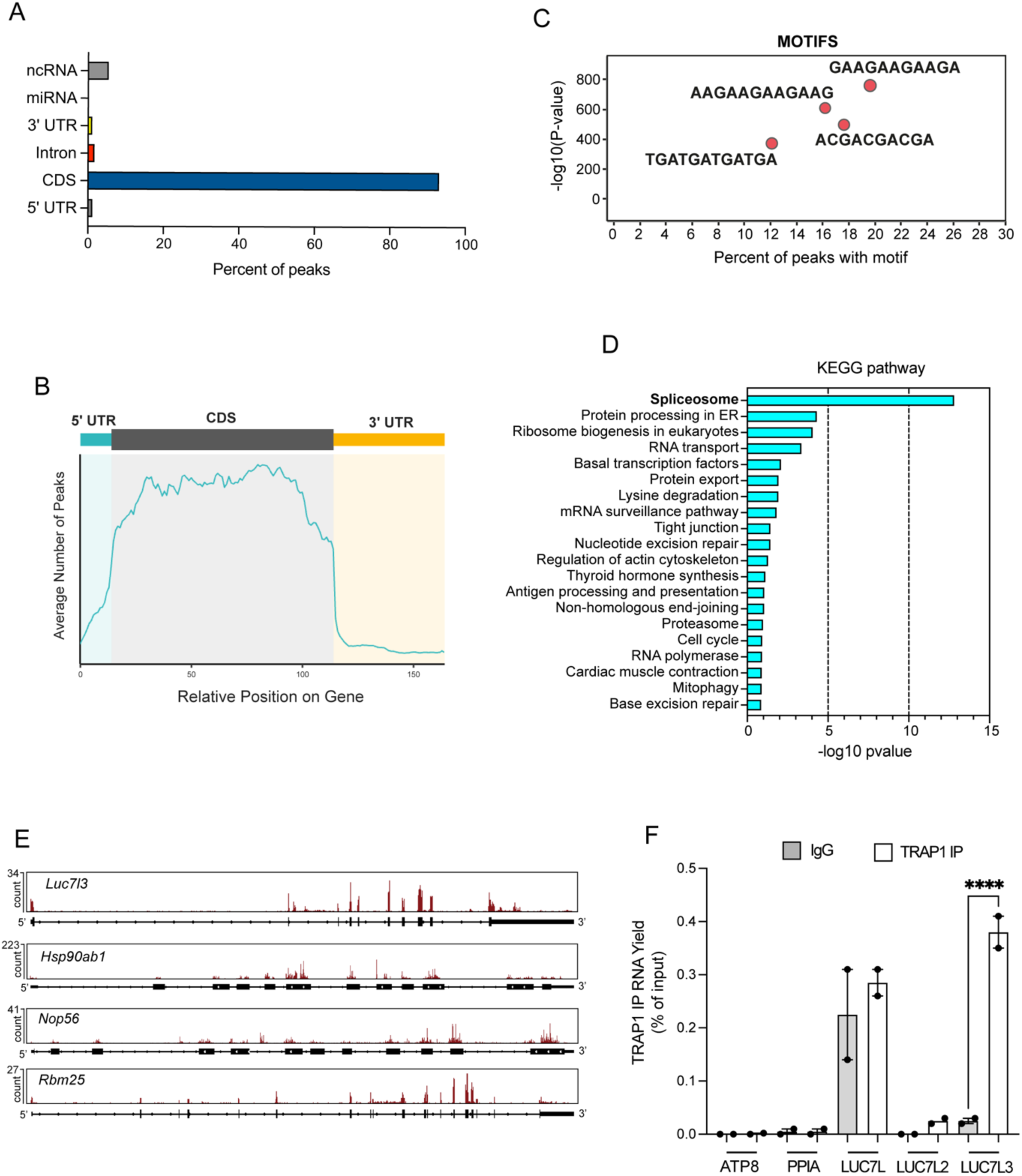
TRAP1 only binds cytosolic transcripts. (**A**) TRAP1 eCLIP density in different transcript regions. (**B**) Meta-analysis of TRAP1 binding around the CDS. (**C**) TRAP1 main binding motifs in PEA1 cells. (**D**) Pathway enrichment analysis of PEA1 reproducible peaks. (**E**) TRAP1 eCLIP profiles for selected genes. (**F**) Formaldehyde crosslinked RNA-IP (RIP) of TRAP1 and control IgG from PEA1 cells. qRT-PCR for LUC7L3 and four control genes. Percentage of immunoprecipitated input is shown for both TRAP1 IP and control IgG. LUC7L, LUC7L, ATP8 and PPIA were used as negative controls. Statistically significant differences were detected using two-way ANOVA and Bonferroni-correction for multiple comparison testing (n = 2). Statistical significance is represented as follows: *p < 0.05, **p < 0.01, and ***p < 0.001. Error bars represent SEM.

### TRAP1 promotes translation of Luc7l3 mRNA

Next, we investigated the effect of TRAP1 binding to *Luc7l3* transcript. First, we performed WB analyses to assess LUC7L3 protein expression levels upon siRNA-mediated TRAP1 silencing in PEA1 and human colorectal carcinoma HCT116 cells, and in HeLa cell clones, where TRAP1 knockdown can be achieved upon tetracycline induction of a TRAP1-directed shRNA. Our data showed that LUC7L3 levels significantly decrease in the absence of TRAP1 only in PEA1 cells (Figure 4A – left panel), as supported by the relative quantification (Figure 4A – right panel), in line with the hypothesis of context-specific functions of TRAP1, typical of several RBPs (Khoroshkin et al., 2024). We performed polysome profiling followed by qPCR analysis on polysomal fractions in PEA1 cells to shed further light on the regulation exerted by TRAP1. Comparison of the polysome profiles from control (siCTRL) and TRAP1 knock-down (siTRAP1) cells revealed no differences between the two conditions (Figure 4B), as confirmed by evaluation of the polysome/monosome ratio (Figure 4C), suggesting that, at least in PEA1 cells, TRAP1 might exert a translational regulation on a restricted group of selective targets. Importantly, analysis of polysomal fractions by qPCR highlighted a significant shift of *Luc7l3* transcript from highly translating polysomes to monosomes in the absence of TRAP1 (Figure 4D – upper panel). This shift is specific for the target, since the housekeeping gene *Ppia* did not follow the same pattern (Figure 4D – lower panel). We also analyzed the steady-state levels of *Luc7l3* mRNA by qPCR to exclude a transcriptional regulation. Indeed, as shown in Figure 4E, TRAP1 silencing did not affect *Luc7l3* mRNA levels in any of the tested cell lines. Altogether, these results indicate that TRAP1 binds *Luc7l3* to promote its translation.

**Figure 4:**
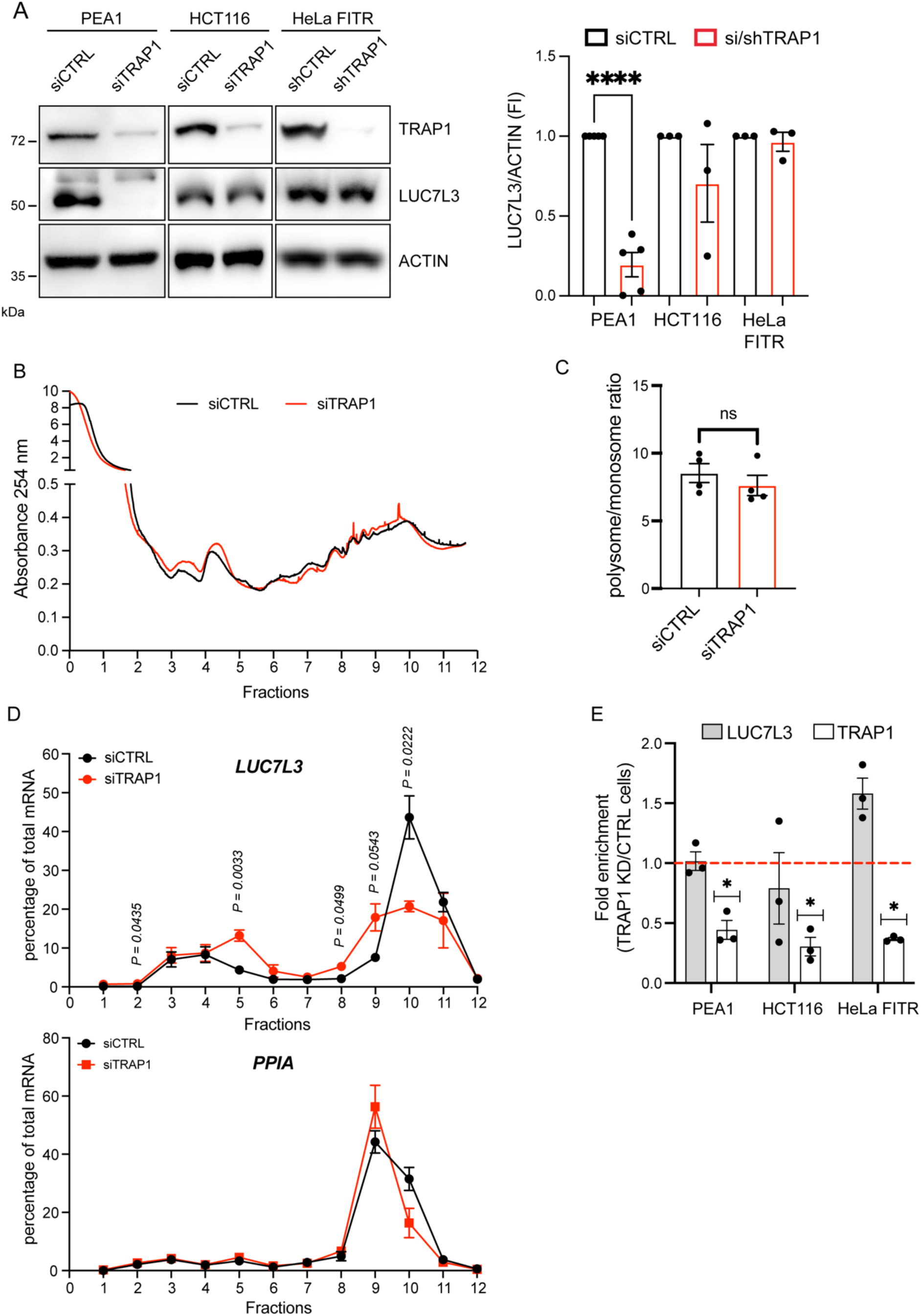
Post-transcriptional regulation of LUC7L3 by TRAP1. (**A**) Assessment of LUC7L3 levels upon TRAP1 modulation. Left, WB analysis of cell extracts upon 72 hrs of silencing using TRAP1-directed siRNA (PEA1 and HCT116) or TRAP1-directed shRNA induction using tetracycline (HeLa FITR). Right, relative densitometric analysis calculated by assuming protein levels of the control equal 1. Statistically significant differences were detected using two-tailed paired Student *t* test (PEA1 n = 5, HCT116 n = 3, HeLa n = 3). Statistical significance is represented as follows: *p < 0.05, **p < 0.01, and ***p < 0.001. Error bars represent SEM. (**B**) Polysome profiling absorbance, measured at 254 nm, of PEA1 cell extracts, from control and siTRAP1 cells. (**C**) Quantification of the polysome/monosome area ratio in control and siTRAP1 profiles as reported in panel A (n = 4). Statistically significant differences were detected using two-tailed unpaired Student *t* test. Error bars represent SEM. (**D**) Percentages of indicated mRNA distributed in sucrose gradient fractions of control and siTRAP1 cells (n = 3). Data are reported as mean ± SEM. Individual P-values are indicated according to two-way ANOVA test. (**E**) LUC7L3 mRNA levels do not decrease after TRAP1 depletion. Transcript levels were assessed by qRT-PCR, corrected for PPIA, and normalized to the levels in shCTRL cells (red line; n = 3). Significance was assessed by one-sample t test. Statistical significance is represented as follows: *p < 0.05, **p < 0.01, and ***p < 0.001. Error bars represent SEM.

### The TRAP1-LUC7L3 axis influences cell proliferation

Besides its well-known role in RNA metabolism through modulation of the splicing process, an essential contribution of LUC7L3 in cell proliferation was recently described (Zhang et al., 2024). We asked whether TRAP1 binding to *Luc7l3* transcript, and its consequent translational regulation, could participate in this process. We performed two in vitro assays in PEA1 cells to test this hypothesis. On one hand, we monitored cell proliferation over four days. To this aim, we knocked down TRAP1 from PEA1 for 72 hours, when the effect of the siRNA reached its peak (data not shown). Then, cells were seeded, and every 24 hours cell proliferation was analyzed using resazurin as a measure of viability. Our data showed that TRAP1 silencing significantly reduces cell proliferation (Figure 5A). On the other hand, we evaluated the ability of cells to grow in isolation by clonogenicity assay. In this case, cells were seeded at low density and monitored until the appearance of distinguished colonies (10-15 days), which were then fixed and stained using crystal violet. We measured the area covered by the colonies at the endpoint. Our results revealed a significant reduction in the ability of cells to form colonies in the absence of TRAP1 (Figure 5B). We also evaluated TRAP1 impact on cell proliferation in PEA1 cells upon stable shRNA-mediated depletion of TRAP1, which confirmed results obtained with the transient silencing (Supplementary Figure 2A-B). LUC7L3 depletion led to similar results (Figure 5C-D). To support the causal link in the TRAP1-LUC7L3 axis and the modulation of cell proliferation, we performed rescue experiments. PEA1 shTRAP1 cells were transfected with wild-type TRAP1 or LUC7L3 expression constructs and their ability to revert the observed phenotype was assessed. Significantly, the effect was partially reverted by expression of either TRAP1 itself or LUC7L3 (Figure 5E-F). These results indicate that, along with additional mechanisms yet to be described, TRAP1 impact on cell proliferation involves LUC7L3 translational regulation. Finally, to evaluate whether TRAP1 and LUC7L3 correlation is conserved in patients, we performed immunohistochemical analyses in HGSOC tissue. This analysis showed that TRAP1 and LUC7L3 are both expressed at high levels in HGSOC (Figure 5G), supporting the potential role of this axis in ovarian cancer progression.

**Figure 5:**
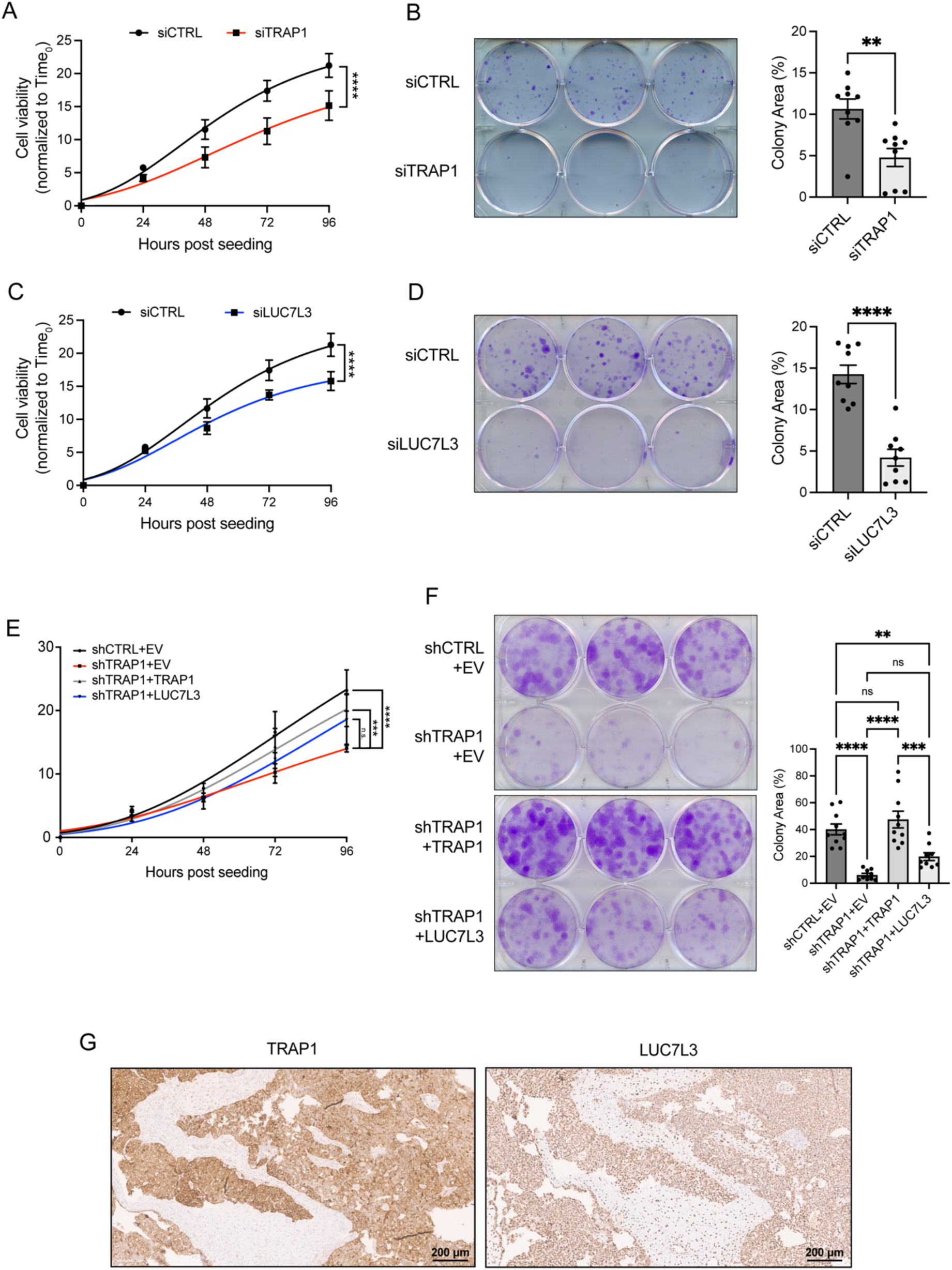
TRAP1 promotes cell proliferation through LUC7L3 regulation. (**A-C**) Effects of TRAP1 or LUC7L3 depletion in proliferation of PEA1 cells. TRAP1 or LUC7L3 were depleted for 72 hrs. Then, siCTRL and siTRAP1/siLUC7L3 cells were seeded in 96-well and cell proliferation was monitored every 24 hrs using Almarblue assay. Significance was assessed by Non-linear fit regression analysis (n = 7). Statistical significance is represented as follows: *p < 0.05, **p < 0.01, and ***p < 0.001. Error bars represent SEM. (**B-D**) Effects of TRAP1 or LUC7L3 depletion in clonogenicity of PEA1 cells. TRAP1 or LUC7L3 were depleted for 72 hrs. Then, siCTRL and siTRAP1/siLUC7L3 cells were seeded in 6-well and cultured for 10-14 days, until the appearance of visible colonies. At the endpoint, cells were fixed and stained with 25% methanol, 0.5% crystal violet. The area covered by the colonies was measured using the ColonyArea ImageJ plug-in. Significance was assessed by unpaired Student’s t test (n = 3). Statistical significance is represented as follows: *p < 0.05, **p < 0.01, and ***p < 0.001. Error bars represent SEM. (**E-F**) PEA1 cells expressing shCTRL or shTRAP1 were transfected with Flag-tagged LUC7L3 or Flag-HA-tagged TRAP1 constructs. 24 hrs post-transfection cells were seeded and assessed for proliferation (n = 5) and clonogenic (n = 3) assays as in (**A**), (**B**), (**C**), (**D**). Statistical significance is represented as follows: *p < 0.05, **p < 0.01, and ***p < 0.001. Error bars represent SEM. (**G**) Immunohistochemical analysis of HGSOC tissue. TRAP1: cytoplasmic staining, please note the focal membranous reinforce. Stromal cells are negative. LUC7L3: nuclear staining. Stromal cells are positive. Images were taken at x200 magnification.

## DISCUSSION

In this study, we provide the first demonstration that the molecular chaperone TRAP1 is an RBP. We validated the RNA binding of TRAP1 in living cells using both protein and RNA-centric complementary approaches. The lack of classical RBDs within TRAP1 and its RNA-unrelated chaperone activity leads us to propose that this protein is an unconventional RBP (Hentze et al., 2018).

Despite its prevalent mitochondrial localization, we reported that TRAP1 partially localizes on the outer face of the endoplasmic reticulum (Amoroso et al., 2012; Avolio et al., 2023). TRAP1 extra-mitochondrial localization is functionally associated with a well-established role in the regulation of protein synthesis, through the binding to both ribosomes and translation factors (Avolio et al., 2023). While TRAP1 globally affects mRNA translation in some settings, it appears to regulate a restricted subset of transcripts in others (Avolio et al., 2023; Matassa et al., 2014). Moreover, the involvement of TRAP1 in protein synthesis is not limited to the cytosol, in fact, TRAP1 is also associated with the mitochondrial translation apparatus, modulating mitochondrial translation (Avolio et al., 2023). The synchronous coordination of these two processes is essential to ensure proper translocation of nuclear-encoded OXPHOS subunits inside mitochondria and assembly of the ETC complexes (Couvillion et al., 2016). Our recent findings align with the hypothesis that TRAP1 could behave as a mammalian chaperone linked to protein synthesis (CLIPS), which have been shown to associate with ribosomes, binding and processing nascent peptides as they emerge from the ribosome exit tunnel (Que et al., 2024). Based on previous reports and given the identification of TRAP1 in the RNA-interactome of several cell lines (Castello et al., 2012; Mullari et al., 2017a; Perez-Perri et al., 2018; Queiroz et al., 2019a; Trendel et al., 2019; Urdaneta et al., 2019), we hypothesized that TRAP1 moonlighting functions in the context of mRNA translation and regulation of cell metabolism involves binding to RNA. OOPS data shown in this article support this hypothesis and show that RNA-binding is regulated following exposure to stress (amino acid deprivation or translation elongation inhibition).

Aiming to further characterize TRAP1 as an RBP, we found that both the ATPase and the ribosomal S5-D2 domains can bind RNA. The former domain has been predicted to be responsible for the RNA-binding properties of HSP90 (Jin et al., 2023b); yet, while this prediction was based on machine-learning-based algorithm no experimental data validated such a prediction. The latter can be involved in both DNA and RNA interactions and, therefore, it is not listed among the canonical RBDs. TRAP1 seems to contact RNA through cooperation between multiple regions, which is expected for some RBPs (Corley et al., 2020).

For a more comprehensive understanding of TRAP1 role as RBP, we identified TRAP1 RNA interactors and obtained two important pieces of information. First, 95% of interactors are protein-coding genes; second, TRAP1 is mainly bound to the CDS of its targets. Regulatory functions of RBP binding to the CDS have been less explored than those associated with binding to the 5’- and 3’-UTRs, for long considered the central regulatory regions mainly responsible for post-transcriptional gene regulation, or to introns, typical of many splicing factors. Nevertheless, a recent study highlighted the impact of RBP binding to CDS in regulating translational efficiency and mRNA stability (Grzybowska & Wakula, 2021). Taken together, these data are in excellent agreement with a prominent role of TRAP1 in the translational regulation of its RNA targets.

Intriguingly, the binding of TRAP1 is exclusive to cytosolic transcripts, although our OOPS data show that TRAP1 RNA binding should be conserved across its different localizations, including mitochondria. In line with our results, recent findings attributed a highly specific RNA-binding function to mitochondrial TRAP1 during heart failure (Liu et al., 2025). In normal conditions, TRAP1 interacts with mitochondria-encoded circRNAs (mecciRNAs), which facilitate TRAP1 interaction with CypD and closing of the mitochondria permeability transition pore (mPTP); whereas, during heart failure, the levels of SUPV3L1/ELAC2 complex increase and mecciRNA are rapidly degraded, thus favoring the opening of the mPTP and release of mitochondrial reactive oxygen species responsible for the deleterious effects (Liu et al., 2025). Considering that the identification of circular RNA requires tailored bioinformatic analyses, whether TRAP1 binds to these types of RNAs in mitochondria of PEA1 cells remains to be elucidated.

One of the most exciting results of this study is that most transcripts bound by TRAP1 encode splicing-related factors. Pre-mRNA splicing is a major mechanism contributing to proteomic diversity in higher organisms (Rogalska et al., 2023). Almost every process inside the cell can be regulated through alternative splicing, which leads to the production of distinct protein isoforms that can show highly different interactions, localizations, and functions (Yang et al., 2016). Dysregulation of splicing events is typically found in cancer cells and can contribute to every hallmark of cancer (Bonnal et al., 2020). For this reason, alternative splicing events and splicing factors are considered effective biomarkers for use in diagnosis, prognosis, and therapy (Bonnal et al., 2020). In this context, the correlation between alternative splicing events and prognosis has been explored in several cancer types, including ovarian carcinoma (Li et al., 2017; Lin et al., 2019; Zhu et al., 2018). A recent study aiming at constructing an independent prognostic signature for predicting ovarian cancer patients’ survival outcomes proposed LUC7L3 as one of the splicing factors responsible for ovarian cancer development (Ouyang et al., 2021). Based on our eCLIP data and the identification of LUC7L3 transcript as TRAP1 RNA ligand, it can be speculated that binding of TRAP1 to splicing factor-encoding transcripts could serve to modulate their expression levels, which, in turn, rewire splicing programs responsible for the regulation of cancer cell traits.

Together with LUC7L and LUC7L2, LUC7L3 is a paralog of yeast Luc7p, which plays a major role in pre-mRNA splicing (Puig et al., 2007). All three proteins bind snRNAs with enrichment at the 5′ ends of U1 and U11 snRNAs and show distinct characteristics in terms of protein interactors and RNA ligand binding preferences (Daniels et al., 2021b). As proof of concept, we confirmed TRAP1-specific binding only to *Luc7l3* transcript and demonstrated that it promotes *Luc7l3* mRNA engagement by actively translating ribosomes and translation. Recent findings demonstrate an important role of LUC7L3 in cell proliferation (Zhang et al., 2024). Notably, our rescue experiments also show that TRAP1 translational regulation of LUC7L3 impacts cell proliferation, in perfect agreement with the above-mentioned study (Zhang et al., 2024). The LUC7L3 protein prevents R-loop accumulation and promotes spindle assembly by facilitating the translation of spindle-related genes, thus ensuring genome stability (Zhang et al., 2024). However, whether this mechanism is conserved in ovarian cancer cells needs further elucidation.

In summary, this study opens a new scenario on the molecular chaperone TRAP1. We show that TRAP1 behaves as an unconventional RBP binding splicing-related factors among its RNA ligands and regulating their translation with consequences for proliferation, opening new avenues to understand ovarian cancer progression.

## METHODS

### Cell culture

Human HCT116 colon carcinoma cells, human cervical carcinoma HeLa cells, and human ovarian carcinoma PEA1 cells were purchased from American Type Culture Collection (ATCC) and cultured in McCoy’s 5A medium (HCT116), DMEM medium (HeLa) and RPMI 1640 medium (PEA1) at 37°C, 5% CO_2_. Culturing media contain 10% fetal bovine serum, 1.5 mmol/L glutamine. The HeLa Flp-In T-REx (FITR) cell line was kindly provided by Dr. Matthias Gromeier (Duke University Medical Center). Generation of the HeLa FITR stable cell lines expressing the GFP- or FLAG-fusion proteins was performed as described in the manufacturer’s protocol (FITR; Invitrogen). HeLa FITR cells were cultured in DMEM supplemented with 10% fetal bovine serum, 1.5 mmol/L glutamine, and appropriate selective antibiotics.

### Constructs and cloning

For TRAP1-eGFP, TRAP1-Flag, TRAP1-.59-Flag, ATPase-Flag, S5-D2-Flag, C-term-Flag plasmids generation, PEA1 cDNA library, eGFP, and FLAG-HA plasmids were used as templates for fusion PCR. The resulting chimeric cDNAs were cloned into pCDNA5/FRT/TO. The same approach was used to generate vectors tagged with Flag-HA containing the MTS sequence upstream of each TRAP1 domain. Similarly, LUC7L3 sequence was cloned into pcDNA3.1-3xFlag.

### Site-Directed Mutagenesis

To generate TRAP1 deletion mutants, a polymerase chain reaction (PCR) amplification was performed using the TRAP1-FLAG construct as a template. Specific primer pairs were designed to amplify nearly the entire plasmid, excluding the region of interest. Subsequently, the Q5® Site-Directed Mutagenesis Kit (New England Biolabs. E0554S) was used to carry out a combined phosphorylation, ligation, and digestion reaction (KLD) of the template plasmid in a single step. The reaction product was transformed into NEB 5-alpha Competent cells, using 50 μl of bacteria for 5 μl of KLD mix. Following transformation, plasmid DNA were isolated, purified, and subjected to Sanger Sequencing to verify the deletion of the region of interest. The primers used are listed below:

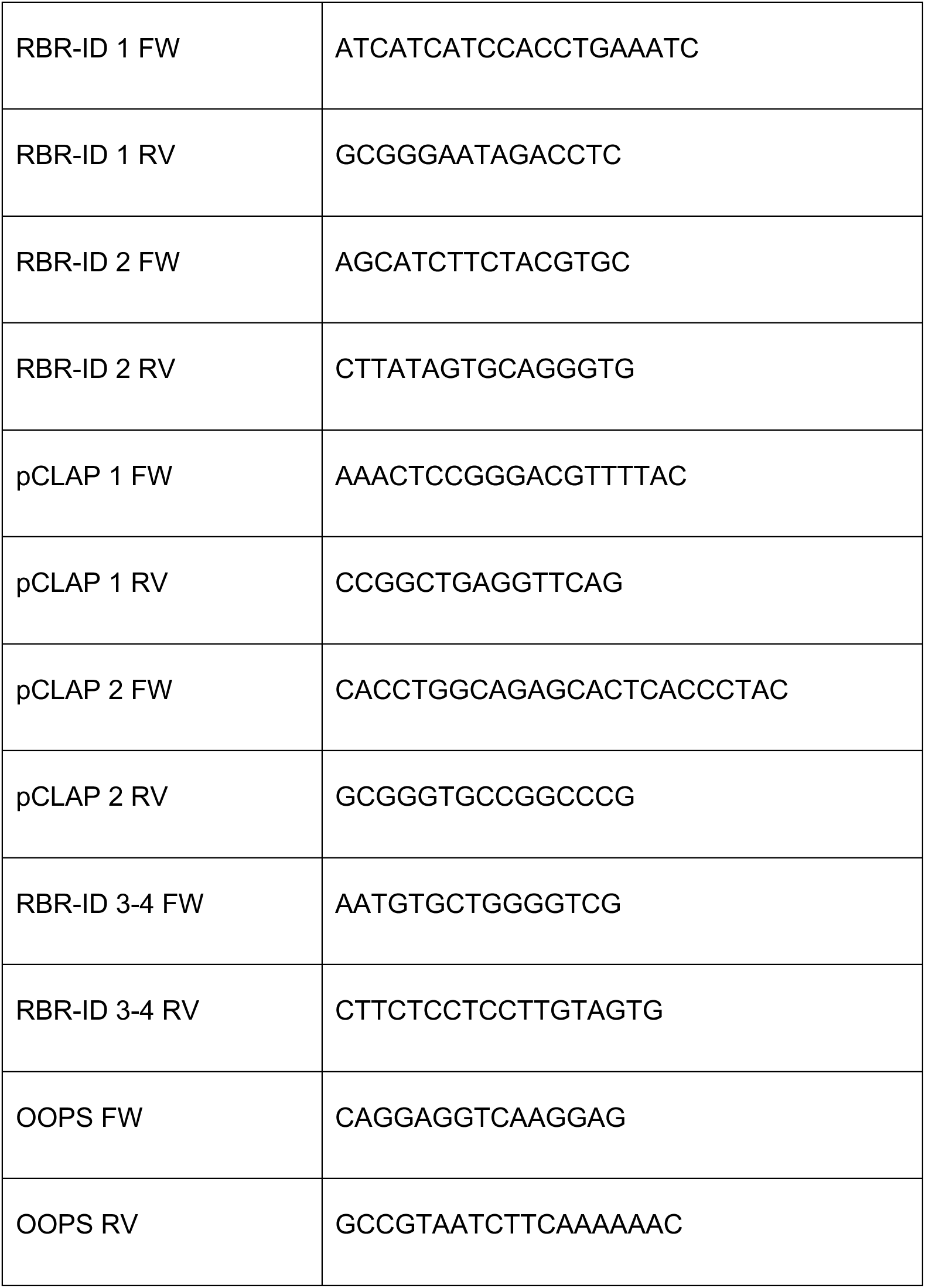

### Viral transduction

Phoenix-Ampho cells were transfected via calcium phosphate protocol to produce viral particles. In detail, 2.5 million cells were seeded in 10 cm plates and the following day they were transfected with a solution containing 437.5 μL of DNA (10 μg of transfer plasmid, i.e. shCTRL-GFP or shTRAP1-GFP), 62.5 μL CaCl_2_ 2M and 500 μL of HEPES-buffered saline solution (HBS:140.5 mM NaCl, 50 mM HEPES and 1.5 mM Na2HPO4 pH 7.12), that was added both dropwise and while vortexing at 1’400 rpm. The plates were then placed at 37°C for 7 hrs to allow the entrance of the plasmid in the cells and, to avoid cellular sufferance, the Phoenix-Ampho cells freshly transfected were then grown in DMEM high glucose, (Gibco, 31966021) supplemented with 10% heat-inactivated FBS (Gibco, 10270106) and 1% penicillin-streptomycin (Gibco, 15070063).

One and two days after transfection, media was collected, filtered (4.5 μm) and supplemented with polybrene diluted 1:1’000. Fresh infections were performed with PEA1 cells at 85% of confluency in 6-well plates, followed by selection with 1 μg/mL puromycin for three days. Finally, to ensure the selection of the stable clones, the PEA1 were sorted through FACS (Fluorescence-activated cell sorting).

### Transfection procedures

For transient silencing of TRAP1 and LUC7L3, we used ON-TARGETplus siRNA (Dharmacon) and performed reverse transfections. Briefly, 2.5 μl of siRNAs (20 μM) were mixed with 2 μl of Dharmafect 1 transfection reagent (Dharmafect, T-2001-03) in serum-free RPMI media, added to 6-well plates and incubated at RT for 30 min. Cells were trypsinized and 80’000 PEA1 cells per well were added. After 72 hrs, cells were trypsinized to start cell culture assays. For clonogenic assays, cells were re-transfected with siRNA every 3 days. In this case, forward transfections were carried out with 1 μl of siRNAs (20 μM) and 0.5 μl of Dharmafect 1 transfection reagent.

For transient plasmid DNA transfection in HeLa cells, Lipofectamine™ 3000 transfection reagent (Thermo Fisher Scientific, L3000015) was used. 400’000 PEA1 cells were plated in p60 mm plates and transfected 24 hrs later. On the day of transfection, two separate mixes were prepared: one containing 4 μg of the plasmid of interest and 8 μl of P300 reagent in serum-free RPMI media; another one, containing 7 μl of Lipofectamine 3000 in serum-free RPMI media. The two mixes were combined and incubated for 15 min at RT before being added to the cells.

### Protein extract preparation and immunoblotting

Equal amounts of protein from cell lysates were subjected to SDS-PAGE and transferred to a PVDF membrane (Merck, IEVH85R). Membranes were blocked in a 5% milk TBS-tween solution for 1 hr at room temperature. The following antibodies were used: anti-HA (Santa Cruz Biotechnology, sc-805 – 1/1000), anti-DDX6 (Novus Biologicals, NB200-191 – 1/1000), anti-H3 (Santa Cruz Biotechnology, sc-517576 – 1/5000), anti-hnRNPC (Proteintech, 11760-1-AP – 1/5000), anti-GFP (Santa Cruz Biotechnology, sc-81045 – 1/500), anti-TRAP1 (Santa Cruz Biotechnology, sc-13557 – 1/1000), anti-LUC7L3 (Proteintech, 14504-1-AP – 1/500), anti-ACTIN (Santa Cruz Biotechnology, sc-47778 – 1/1000). All primary antibodies were incubated overnight at 4°C. Anti-mouse (Bethyl laboratories, A90-137P) and anti-rabbit (Bethyl laboratories, A120-108P) antibodies were used at a 1:20000 dilution and incubated for 1 hr at room temperature. Images were acquired with a Chemidoc MP imaging system (Bio-Rad), and where indicated, protein levels were quantified by densitometric analysis using the software ImageJ (Schneider et al. 2012).

### Cross-linking and immunoprecipitation (CLIP) assay

HeLa FITR TRAP1-Flag-HA cells were cross-linked on ice at 0.30 J/cm^2^ with UV light at 254 nM. Immediately after irradiation, cells were lysed in 1mL lysis buffer (50 mM Tris-HCl, pH 7.4; 100 mM NaCl; 1% NP-40; 0.1% SDS; 0.5% sodium deoxycholate). Cell extracts were sonicated using a bioruptor (Digenode) for 10 cycles of 30 seconds, level L at 4°C, and then cleared by centrifugation at 10’000 rpm for 10 min at 4°C. RNA was then partially digested with either low (1/200) or high (1/25) RNase I (Ambion, AM2294), as well as 2 ml of Turbo DNase (Thermo Fisher Scientific, AM2238) and incubated for 5 min at 37°C under shaking at 1100 rpm. TRAP1-Flag-HA was then captured by incubation with 20 μl of anti-FLAG^®^ M2 magnetic beads (Merck, M8823) for 2 hrs at 4°C with gentle rotation. Beads were then washed 2x with 900 μl of high-salt buffer (50 mM Tris-HC lpH 7.4, 1 M NaCl, 1 mM EDTA, 1% NP-40, 0.1% SDS, 0.5% sodium deoxycholate), 1x with 900 μl PNK wash buffer (20 mM Tris-HCl pH 7.4, 10 mM MgCl_2_, 0.2% Tween-20) and resuspended in 40 μl of PNK mix (30 μl water, 8 μl 5x PNK pH 6.5 buffer [350 mM Tris-HCl pH 6.5, 50 mM MgCl_2_, 5 mM DTT],1 μl PNK enzyme [New England Biolabs, M0201L], 0.5 μl RNasin [Promega, N2611], 0.5 μl of FastAP alkaline phosphatase [Thermo Fisher Scientific, EF0651]) and incubated for 40 min at 37°C under gentle shaking at 1100 rpm. Beads were washed once with 900 μl of ligation buffer (50 mM Tris-HCl pH 7.5, 10 mM MgCl_2_) and resuspended in 25 μl of ligation mix (6.8 μl water, 2.5 μl 10X ligation buffer, 0.8 μl 100% DMSO, 2.5 μl T4 RNA ligase I– high concentration [New England Biolabs, M0437M], 0.4 μl RNasin [Promega, N2611], 0.5 μl PNK enzyme, 2.5 μl pre-adenylated adaptor L3-App [1 mM] conjugated to an infrared-dye [IRdye-800CW-DBCO (LICOR)], 9 μl 50% PEG8000). Ligation was performed at room temperature for 75 min flicking the tubes every 10 min. Beads were washed twice with high-salt buffer, once with PNK wash buffer, resuspended in 20 μl of loading buffer and run on a 4-12% NuPAGE Bis-Tris gel (Invitrogen, NW04122BOX) in MOPS buffer according to the manufacturer’s instructions. Protein-RNA complexes were transferred to a nitrocellulose membrane. Protein infrared-RNA complexes were visualized at 800 nm with a Chemidoc MP imaging system (Bio-Rad).

For biotin/streptavidin-based CLIP, the Antibody IP Validation Kit (ECLIPSEbio, ECAV-0001) was used according to the manufacturer’s instructions. PEA1 cells were *in vivo* UV-crosslinked as described before. For TRAP1 immunoprecipitation 5 μg of TRAP1 antibody conjugated to 50 μl of protein G Dynabeads (Invitrogen, 10004D) were used.

### RNA Interactome capture for eGFP-fused protein

1×15 cm plate of eGFP-fusion protein expressing cells was induced for 24 hrs (TRAP1) and 16 hrs (eGFP) with 1 μg/ml doxycycline. TRAP1-eGFP and eGFP cells were treated with 100 μM 4-thiouridine overnight and photoactivatable ribonucleoside-enhanced crosslinked (PAR-CL) on ice at 0.60 and 0.30 J cm^−2^ with UV light at 365 nm. Following UV-irradiation the protocol was performed as previously described (Strein et al., 2014).

### Orthogonal Organic Phase Separation (OOPS)

OOPS was performed as described in (ref). Briefly, cells were UV cross-linked on ice at 0.30 J/cm^2^ with UV light at 254 nM. Then, cells were washed twice with PBS and immediately lysed by scraping in TRIzol (Thermo Fisher Scientific, 15596018). The homogenate was transferred to a new tube and incubated at RT for 5 min to dissociate unstable RNA–protein interactions. For biphasic extraction, 200 μL of chloroform (Sigma Aldrich, 319988) were added and samples were centrifuged for 15 min at 12’000 g at 4 °C. The upper, aqueous phase (containing non-cross-linked RNAs) and the lower, organic phase (containing non-cross-linked proteins) were discarded and the interface (containing the protein–RNA adducts) was subjected to two more AGPC phase separation cycles, precipitated by the addition of nine volumes of methanol, and pelleted by centrifugation at 14’000 g, RT for 10 min. For RNA-binding protein analyses, the precipitated interface was resuspended in 100 μL of 100 mM triethylammonium bicarbonate (TEAB) and 1% SDS, sonicated using a bioruptor (Digenode) for 15 cycles of 30 seconds (ON/OFF), level H at 4°C, incubated at 95 °C for 20 min, cooled and digested with 2 µl RNase A/T1 (Thermo Fisher Scientific, EN0551) overnight at 37 °C. The following day, after a final cycle of AGPC phase partitioning, proteins were released and recovered from the organic phase by methanol precipitation and analyzed by western blot.

### eCLIP-seq library preparation

Enhanced crosslinking and immunoprecipitation sequencing (eCLIP) libraries were constructed using the RBP-eCLIP kit (ECLIPSEbio) according to the manufacturer’s instructions. Briefly, PEA1 cells were UV cross-linked on ice at 0.40 J/cm^2^ with UV light at 254 nM, followed by cell lysis and lysate treatment with RNAse I to shear RNA. TRAP1 protein-RNA complexes were immunoprecipitated. An RNA adapter was ligated to the 3′ end of TRAP1-bound RNA. Proteinase K was used to digest proteins before reverse transcription of RNA. cDNA was ligated to the 3′ end of single-stranded DNA adapters and amplified. The final libraries were sequenced by Illumina NextSeq 550 (75 bp single-end reads) and 30 million reads were collected for each sample.

### eCLIP-seq data processing

Samples were processed with Eclipsebio’s proprietary analysis pipeline (v1). UMIs were pruned from read sequences using umi_tools (v1.1.1). Next, 3’ adapters were trimmed from reads using cutadapt (v3.2). Reads were then mapped to a custom database of repetitive elements and rRNA sequences. All nonrepeat mapped reads were mapped to the genome GRCh38 (hg38) using STAR (v2.7.7a). PCR duplicates were removed using umi_tools (v1.1.1). RBP-eCLIP peaks were identified within IP samples using the peak caller CLIPper (v2.0.1). For each peak, IP versus input fold enrichments and p-values were calculated. Peaks were annotated using transcript information from GENCODE v41 (GRCh38.p13) with the following priority hierarchy to define the final annotation of overlapping features: protein coding transcript (CDS, UTRs, intron), followed by non-coding transcripts (exon, intron).

### Formaldehyde cross linked RNA Immunoprecipitation (fRIP)

fRIP was performed as described in (Chatterjee et al., 2024)with minor modifications. For antibody-beads conjugation, 30 µl of Protein G magnetic bead slurry (Invitrogen, 10004D) and 5 µg antibodies were incubated overnight at 4°C with gentle rotation. For each IP reaction, 3 mg lysate (in 300 µl of RIPA) was used. RNA was extracted with TRIzol and purified with ZYMO RNA Clean & Concentrator-5 kit (Zymo, #R1016). cDNA was prepared from whole purified RNA with OneScript® Plus cDNA Synthesis Kit (Abm, G236) using random primers.

### RNA Extraction and RT-qPCR

Total RNA extraction procedures were performed by using TRIzol, following the manufacturer’s instructions. For first-strand synthesis of cDNA, 1 µg of RNA was used in a 20 µL reaction mixture by using the All-In-One 5X RT MasterMix with gDNA Removal (Abm, G592). For real-time PCR analysis, 20 ng of cDNA sample was amplified by using the BlasTaq™ 2X qPCR Master Mix (Abm, G891) in a CFX96 Touch Real-Time PCR Detection System (Bio-Rad). The primers used are listed below:

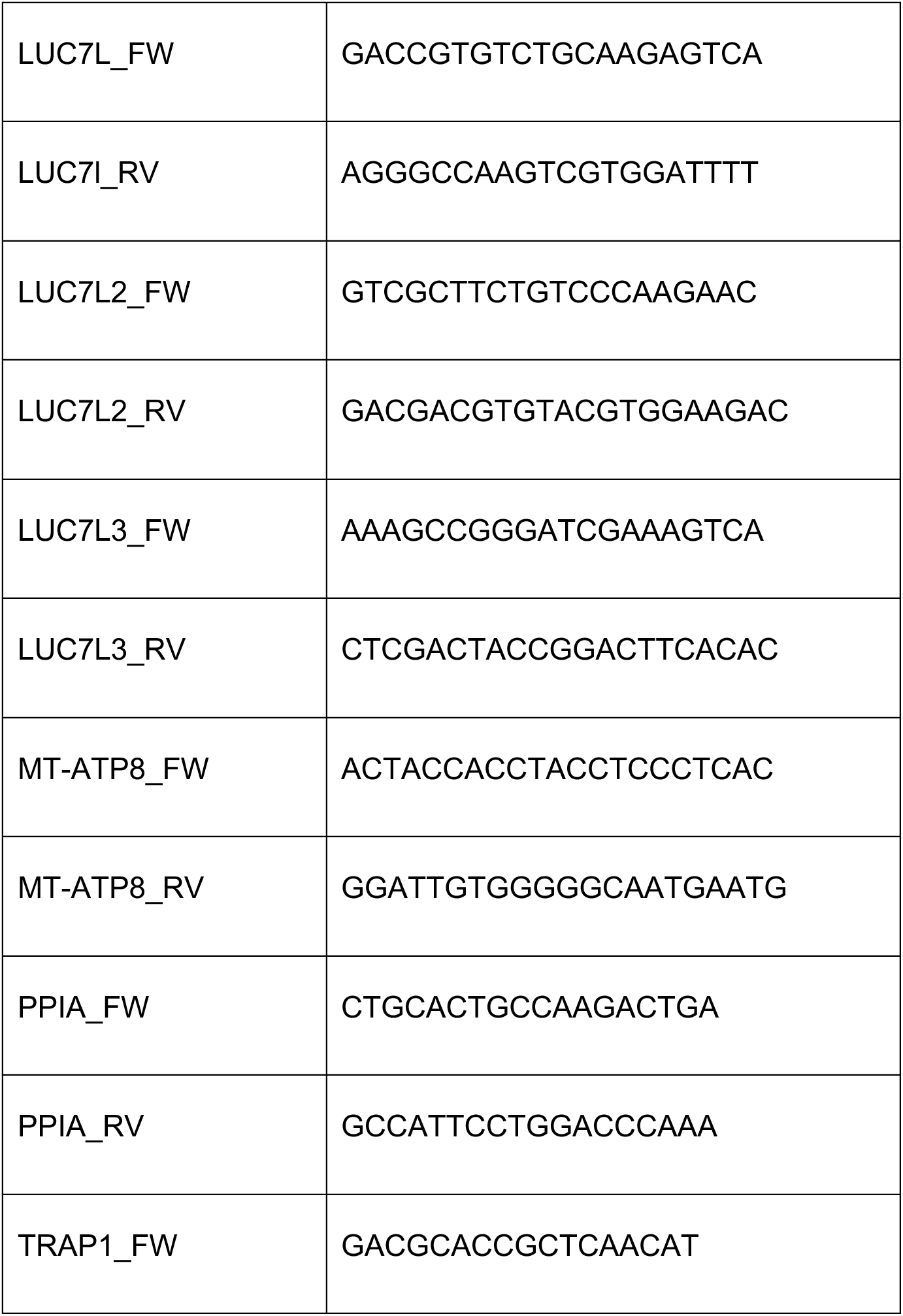
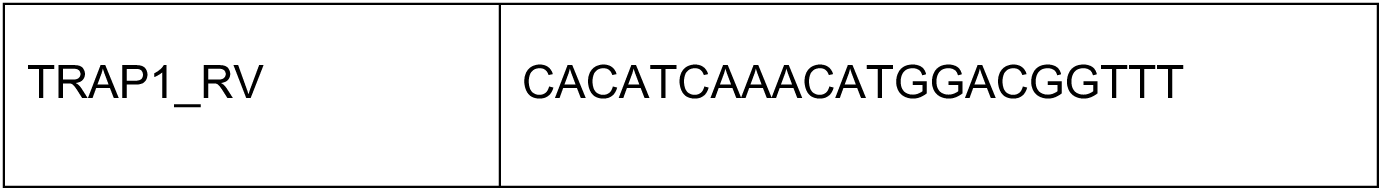

### Polysome Profiling

Following 72 hours of transfection with siCTRL and siTRAP1, PEA1 cells were washed with ice-cold PBS supplemented with 100 µg/mL cycloheximide and resuspended in 500 µL of lysis buffer (10 mM Tris-HCl at pH 7.4, 100 mM KCl, 10 mM MgCl_2_, 1% Triton X-100, 1 mM DTT, 10 U/mL RNaseOUT [Invitrogen, 10777019], 100 µg/mL of cycloheximide). After 10 min of incubation on ice, cell lysate was centrifuged for 10 min at 14’000 rpm at 4°C. The supernatant was collected, and protein concentration was determined using Bradford assay (Bio-rad). Equal amounts of protein were loaded onto a 10%–60% sucrose gradient obtained by adding 6 mL of 10% sucrose over a layer of 6 mL of 60% sucrose prepared in lysis buffer, in a 12-mL Open-Top Thinwall Ultra-Clear Tube (Beckman, 344059). Gradients were prepared using a gradient maker (BioComp). Polysomes were separated by centrifugation at 37’000 rpm for 2 hrs using a Beckmann SW41 rotor. Twelve fractions of 920 µL were collected, and polysomes were monitored by following the absorbance at 254 nm. Following the addition of 50 pg of Luciferase RNA as an internal control to each fraction, RNA extraction was performed by adding TRIzol to the fractions at a 1:1 v/v ratio following the manufacturer’s instructions.

### Analysis of Polysomal-associated mRNA

The same volume of RNA from each fraction was reverse transcribed to cDNA and used for qPCR assays. RNA amounts were quantified, normalized to internal Luc as control, summed across all fractions, analyzed and presented after normalized to their total RNA concentrations.

### Cell viability assays

Following 72 hours of transfection with siTRAP1 or siLUC7L3, PEA1 cells were detached from the plate and seeded in a 96-well plate (1’500 cells/well) in a final volume of 100 µl. At 24, 48, 72, and 96 hrs post-seeding, 10 µl alamarBlue™ HS Cell Viability Reagent (Invitrogen, A50101) were added to the culture medium. After 4-6 hrs incubation at 37°C, fluorescence intensity of resorufin was measured using a SynergyH1 microplate reader (BioTek).

### Clonogenic assays

Following 72 hours of transfection with siTRAP1 or siLUC7L3, PEA1 cells were counted and plated at a low density (1’000 cells per well) in 6-well plates and incubated for 10-14 days. siRNA replenish was performed every 3 days to ensure TRAP1 and LUC7L3 silencing throughout the experiment. Following the incubation period, cells were washed with PBS and stained with 0.5% crystal violet and 25% methanol, then incubated for 1 hr at RT. Subsequently, two additional washes with H_2_O were performed, and cells were left to air dry until complete evaporation. The area covered by the colonies was measured with the plug-in ColonyArea of ImageJ (Guzmán et al., 2014).

### Immunohistochemical analysis

Immunohistochemical staining procedures were carried out on formalin-fixed, paraffin-embedded cell block. Serial tissue sections (4μm-thick) were prepared for immunohistochemical analysis. We used Ventana BenchMark ULTRA IHC/ISH (Ventana Medical Systems, Tucson, AZ) immunostainer, applying the OptiView DAB IHC Detection Kit (OptiView) an indirect, biotin-free system. Anti-TRAP1 (Santa Cruz Biotechnology, sc-13557 - dilution 1:500), was incubated for 60 min at 36C°, while anti-LUC7L3 (Proteintech, 14504-1-AP - dilution 1:20) was incubated for 40 min at 36C°, followed by optiview amplification kit application for 8+8 min. Hematoxilin and bluing reagent included in the immunostainer protocol was perfomed. Two pathologists evaluated the reaction independently.

### Gene ontology

For all gene ontology analyses the enrichR webtools was used (Kuleshov et al., 2016).

### Statistical Analyses

GraphPad Prism 9 (Version 9.3.1) software was used for statistical data analysis. Bar graphs display mean±SEM and all data points. Experimental replicates are mentioned in the figure’s legends. Statistical significance is represented in all figures, as follows: p < 0.05:*, p < 0.01:**, p < 0.001:***, p < 0.0001:****, and p-value of ≥0.05: not significant.

## Data availability

Sequencing data have been submitted to GEO and are available under accession number PRJNA1241711.

## Supplementary data

Supplementary data are available for this article.

## Acknowledgments

We thank Fatima Gebauer for carefully reading the manuscript. We acknowledge the sequencing facility at the Department of Molecular Medicine and Medical Biotechnology, University of Napoli “Federico II,” for the support in eCLIP libraries sequencing.

## Author contributions

Conceptualization. R.A., D.S.M. and F.E. Investigation. S.D.L., R.A., L.P., M.A.A.A., C.M., R.C. Formal data analysis and interpretation. S.D.L., R.A., D.S.M. Project administration. R.A., D.S.M. Mentoring and supervision. R.A. and D.S.M. Writing – original draft. R.A. Writing – review and editing. R.A., D.S.M. and F.E.

