## Supplementary_Figure_1 for "The molecular chaperone TRAP1 promotes translation of *Luc7I3* mRNA to enhance ovarian cancer cell proliferation"

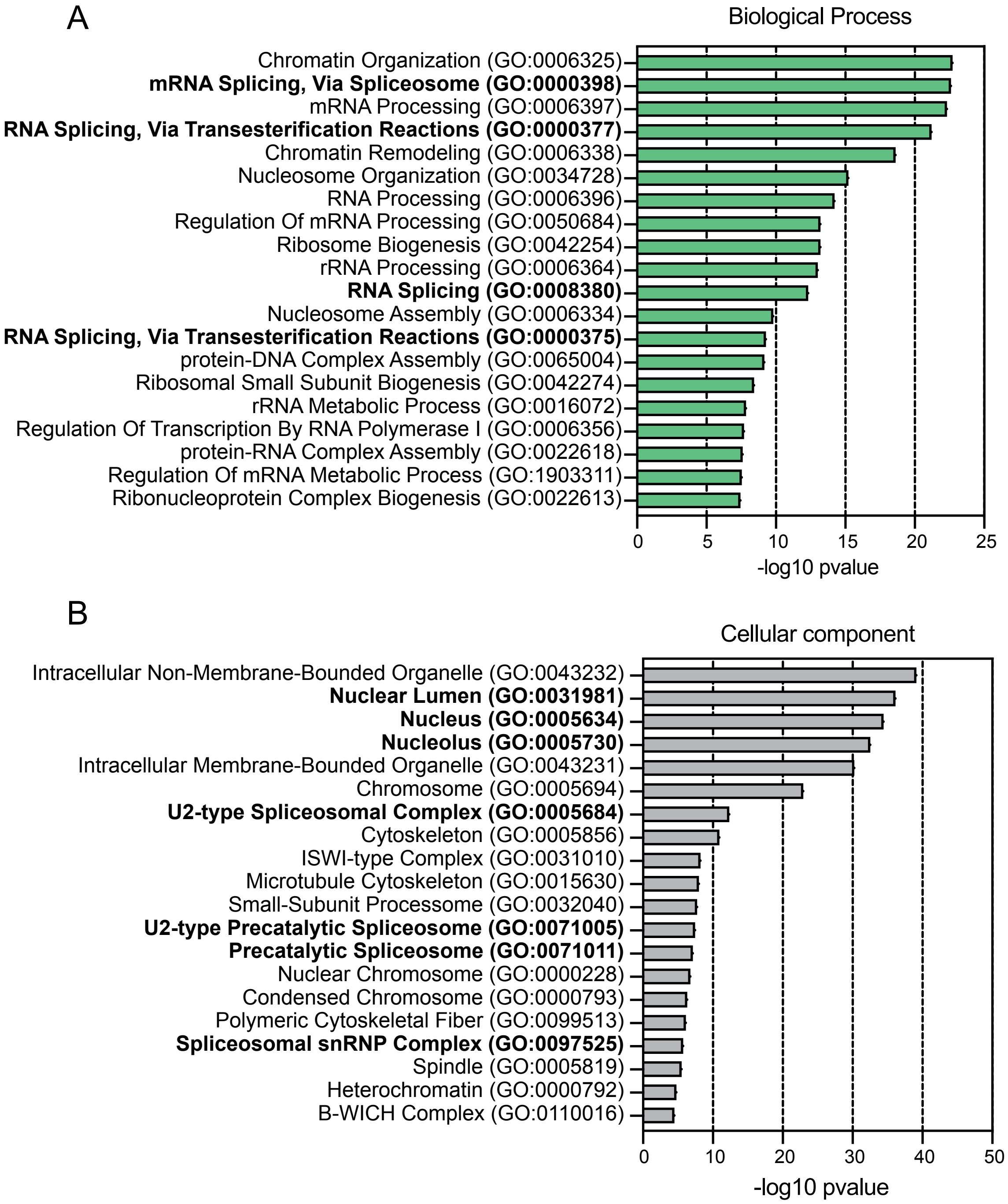


**Supplementary Figure S1**: (**A**) Biological processes enrichment analysis of PEA1 reproducible peaks. (**B**) Cellular component enrichment analysis of PEA1 reproducible peaks.
