## Supplementary_Figure_2 for "The molecular chaperone TRAP1 promotes translation of *Luc7I3* mRNA to enhance ovarian cancer cell proliferation"

A

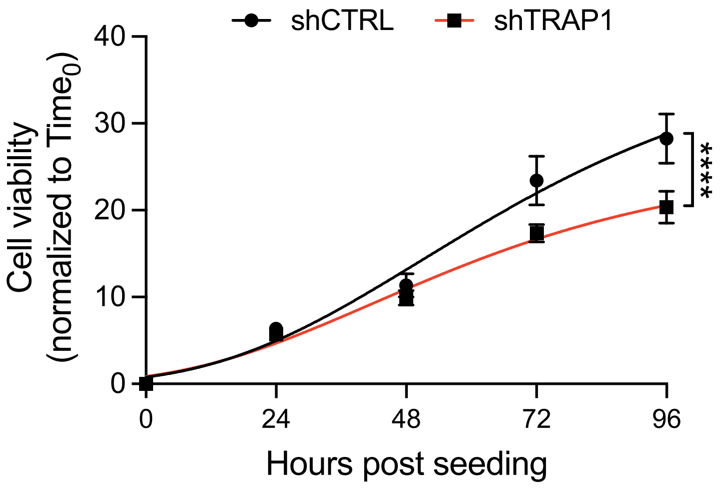


B

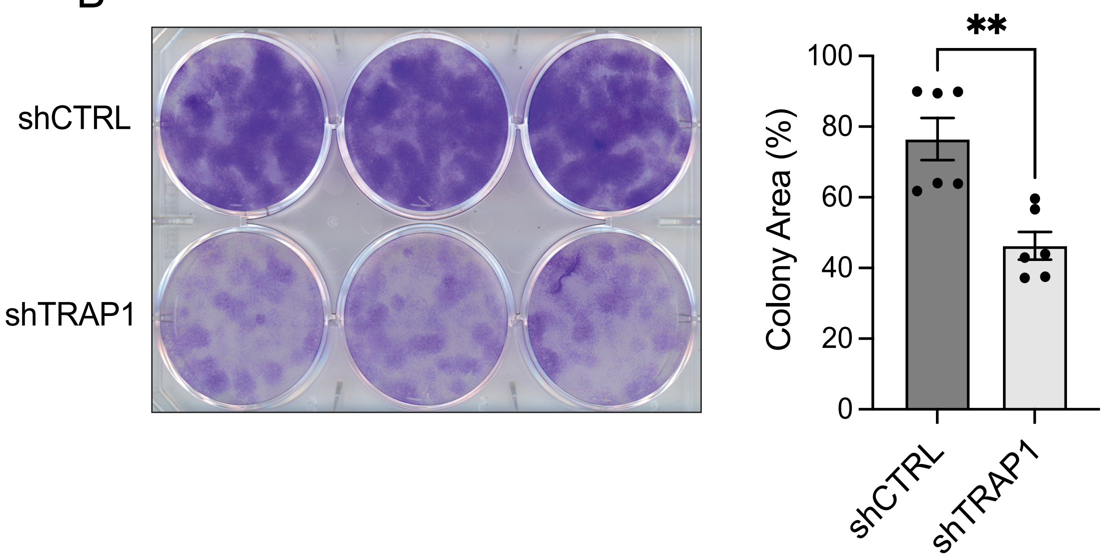


**Supplementary Figure S2**: (**A**) Effects of TRAP1 depletion in proliferation of PEA1 cells. shCTRL and shTRAP1 cells were seeded in 96-well and cell proliferation was monitored every 24 hrs using Almarblue assay. Significance was assessed by Non-linear fit regression analysis (n = 7). Statistical significance is represented as follows: *p < 0.05, **p < 0.01, and ***p < 0.001. Error bars represent SEM. (**B**) Effects of TRAP1 depletion in clonogenicity of PEA1 cells. shCTRL and shTRAP1 cells were seeded in 6-well and cultured for 10-14 days, until the appearance of visible colonies. At the endpoint, cells were fixed and stained with 25% methanol, 0.5% crystal violet. The area covered by the colonies was measured using the ColonyArea ImageJ plug-in. Significance was assessed by unpaired Student's t test (n = 3). Statistical significance is represented as follows: *p < 0.05, **p < 0.01, and ***p < 0.001. Error bars represent SEM.
